# Leveraging existing data to maximise quality and consistency across gene model annotations: a *Fusarium* pan-annotation

**DOI:** 10.1101/2025.03.12.642647

**Authors:** Rowena Hill, Gillian Reynolds, Neil Hall, David Swarbreck

## Abstract

Comparative genomics analyses are frequently used to inform our understanding of how organisms have evolved and how genetics contributes to phenotypic traits. Thanks to the considerable growth in the number of sequenced and assembled genomes, there is increasingly abundant data with which to perform such analyses. However, the appropriate use of genomic data can be highly dependent on genome annotation. Genome annotation is the critical step for providing biological context to genome sequences, but due to the complexity of the task it remains a major bottleneck in analyses. Different methods, and a lack of widely accepted standards, can also result in a great diversity of completeness and accuracy across genome annotations. Accordingly, comparative genomics analyses are susceptible to errors which can be misinterpreted as biological variation, yet efforts to revise and update existing annotations have not kept pace with advances in technology and expanding data resources. We have developed a workflow to utilise existing genome annotations alongside *de novo* gene predictions to improve both the collective consistency and individual quality of genome annotations of a closely related group of genomes. In this work we apply this new workflow on a dataset of 82 genomes from the economically, ecologically and clinically important fungal genus *Fusarium*. We show that both individual as well as collective annotation quality can be improved. The development of reannotation approaches such as we present here will be essential if we are to capitalise on the huge investment that has gone into generating existing genome data.

The *Fusarium* pan-annotation is available from Zenodo at https://zenodo.org/doi/10.5281/zenodo.13829922. Workflow code and sample commands are available from https://github.com/EI-CoreBioinformatics/FusariumPanAnno.

**KEY POINTS:**

- Comparative genomics is a fundamental approach to understand the contributions of genetic features to biological questions.
- To take advantage of existing data, most comparative genomics studies compare gene models produced using a variety of annotation methodologies, which introduces computational bias that can be misinterpreted as biological signal.
- We present a bioinformatics workflow to improve consistency across a set of gene model annotations and minimise computational bias for downstream comparative genomics analyses.
- Reannotation of a previously published *Fusarium* dataset reaffirms the finding that comparing annotations generated from a mix of methodologies can underestimate core genes, overestimate taxon-specific genes and confound patterns of gene presence/absence.

## INTRODUCTION

The start of the sequencing revolution in the latter half of the 20^th^ century [1] has led to an explosion in sequence data, a major focus of which has been genome assemblies. While initially these assemblies, pieced together from short-read data, were highly fragmented and incomplete, the development of long-read sequencing technologies has facilitated an increase in the number of complete, or near-complete, genome assemblies with improved contiguity, accuracy and, in some cases, genomic resolution with haplotype-aware assemblies (e.g. [2]). Major initiatives, such as the Darwin Tree of Life (DTOL) [3] and the European Reference Genome Atlas (ERGA) [4] are capitalising on these gains in sequencing capacity and affordability to expand the pool of genome assemblies available to biologists, which will provide unparalleled insight into the sequences which encode Earth’s diverse lifeforms [5]. However, genomic sequences, lack biological context without an accompanying genome annotation, that decorate the genomic data with biological features. Annotation enables an array of downstream analyses such as comparative genomics and phylogenomics [5]. Accurate gene annotation is also foundational to the development of synthetic biology and bioengineering methods [6].

Given the rapid rate of species loss globally [7], it is critical that we catalogue, analyse and understand as much as possible, as quickly as possible. Yet genome annotation methods are a major bottleneck for genomic analysis [8]. This is due to the underlying computational complexity of the annotation methods, the scale and variety of data that supports annotation, as well as the biological complexity and diversity of genomes across the tree of life. The gold-standard for gene model prediction typically involves a combination of evidence-based transcriptome, protein alignment and gene prediction utilising probabilistic models and integration of extrinsic data, but individual annotation tools/workflows can vary in their specific approach, with different strengths and weaknesses [9,10]. High quality annotation is also dependent on access to bioinformatics expertise and computational resources. The result is that genome annotation is often performed for a small fraction of genomes, with varying completeness and accuracy. Further, the process of re-visiting, correcting and updating genome annotations is performed in only a minority of cases [11], and novel methodologies to improve annotations are rarely taken advantage of for anything but the newest genome sequencing efforts. This mosaic of annotation efforts and lack of widely agreed-upon standards [12] makes comparative genomics efforts fraught with errors which are prone to misinterpretation as biological variation [13–17]. All these factors hamper efforts to characterise and understand the genomic variation which underpins phenotypic traits of interest.

If we are to make best use of the significant time and investment that has gone into producing the huge quantity of existing genomic data, strategies are needed which can minimise the impact of different annotation methodologies across comparative genomics analyses. Here, we demonstrate a holistic approach to improve individual genome annotations, but also increase consistency across annotations as a collective. As a case-study, we used a dataset of 62 taxa in the fungal genus *Fusarium*, which was previously compiled to assess the contribution of gene content to fungal lifestyle in the lineage [18]. *Fusarium* is an ecologically, economically and clinically important genus of globally distributed microfungi [19], for which minor differences in gene presence/absence between strains have previously been correlated with differences in pathogenicity and host specificity [20,21]. We have developed a workflow in which existing genome annotations can be leveraged to generate a collection of gene models which can then be projected across a group of closely related genomes. We complement this projection approach with additional *denovo* gene annotation. This combinatorial process improves both individual genome annotation quality and collective consistency, reducing the technical bias inherent to comparative genomic work which utilises genome annotations produced from differing methods.

## METHODS

### Data acquisition

82 *Fusarium* genomes and 1 *Ilyonectria* genome were obtained from NCBI GenBank and JGI MycoCosm (Supplementary Table S1).

### Reannotation

The pan-reannotation process implements a complementary combination of *de novo* gene predictors and homology-based gene liftovers to maximise the pool from which the best gene model can be selected for each locus, within each genome. An overview of the method can be seen in Fig. 1. The first stage of the process is to generate a set of gene models for each genome. This is achieved by combining novel gene prediction via Galba v1.0.9 [22] and Helixer v0.3.1 [23] and pairwise gene alignments of gene models from all genomes via LiftOff v1.5.1 [24].

**Figure 1.**
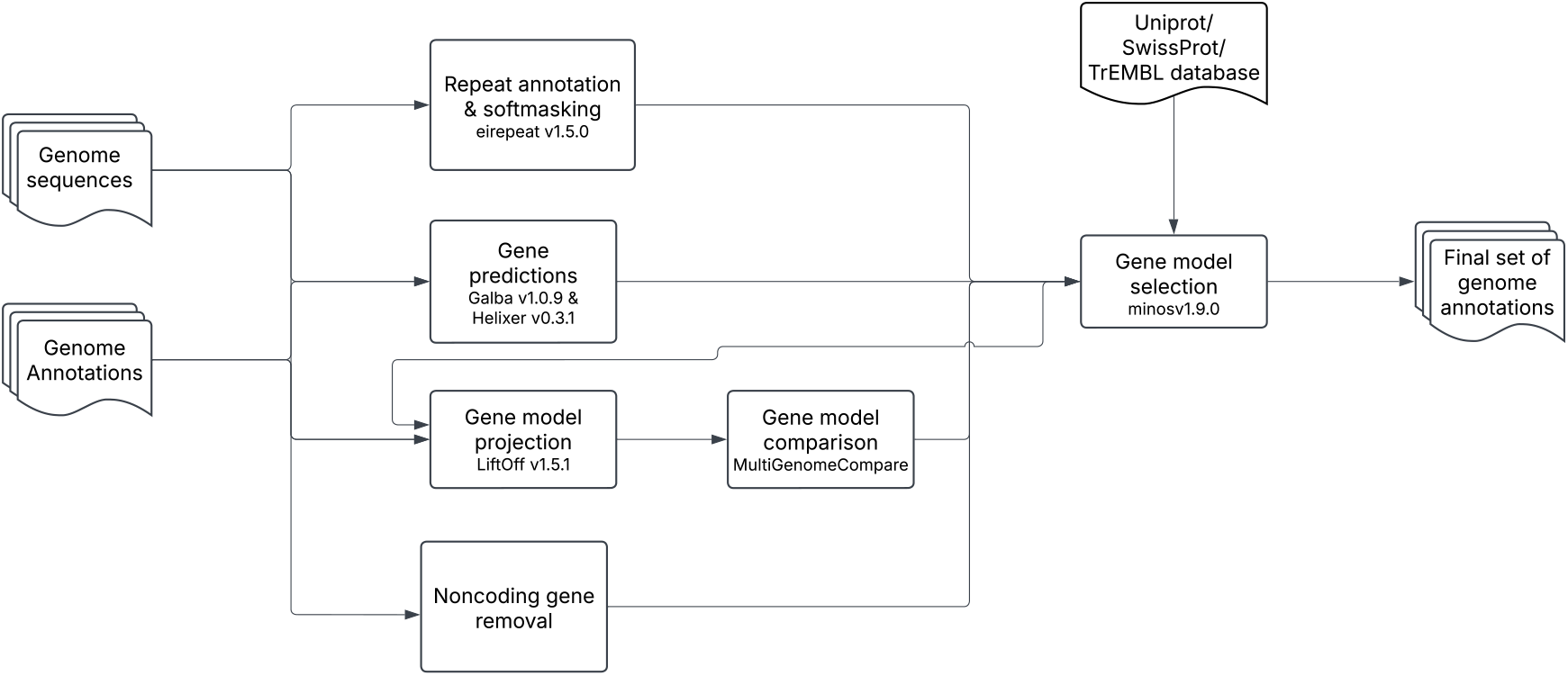
Schematic of reannotation workflow.

The use of both Galba and Helixer optimises *de novo* gene prediction by leveraging the distinct algorithmic approach of each – where Galba uses a classic Hidden Markov Model capable of integrating biological data via spliced protein alignment of closely related species proteomes, Helixer uses a novel Deep Neural Network approach which only requires genome sequence, annotation data and a pre-trained model [23,24]. We utilised Helixer’s pre-trained fungal model (fungi_v0.3_a_0100) available from https://zenodo.org/records/10836346. As Galba requires a set of proteins from multiple closely related species to perform optimally, proteins from all accessions except those excluded on quality grounds (**ΔBUSCO** ≤ 100 as determined via compleasm [25], see description of metric below and ≤ 2000 unknown genes and contamination as determined via OMArk v0.3.0 [26]) were combined. The included and excluded accessions are shown in Supplementary Datasheet S1. The OMAmer database used by OMArk was LUCA.h5 (available from https://omabrowser.org/oma/current/). Proteins were extracted from each assembly via gffread v0.12.7 [27].

To complement novel gene model prediction, existing gene annotations for all genomes within our dataset were leveraged via pairwise projections of gene models via LiftOff v.1.5.1. Successfully projected gene features were then assessed via MultiGenomeCompare (https://github.com/lucventurini/ei-liftover/) with only those models transferred fully with no loss of bases and identical exon/intron structure retained (F1 score of 100 for CDS, Exon and CDS; Junction alongside class code indicating complete intron chain matches; or identical monoexonic transcripts).

The output of the above processes, alongside a filtered set of the original annotations, were fed into minos v.1.9.0 [28] for selection of optimal gene models. Also provided to minos was: (1) a database of *Hypocreales* (taxid 5125) proteins downloaded from Uniprot with any proteins belonging to the *Fusarium* genus (taxid 5506) removed; (2) BUSCO v5.5.0 [29] assessment of original annotations performed using the hypocreales_odb10.2019-11-20 database; (3) extracted proteins from original annotations via gffread v0.12.7; and (4) interspersed-repeat annotations obtained via eirepeat v1.5.0 [30], which integrates RepeatModeler [31] and RepeatMasker [32]. For each genome, gene models were scored and selected via minos from the original annotation, Galba, Helixer and LiftOff projected models based on consistency and extent of supporting data (i.e. the protein and LiftOff datasets).

Following gene model selection via minos, to aid annotation consistency a final round of pairwise liftovers of gene models was perfomed using LiftOff. The projected gene models were then provided as input to minos, along with the first round minos and Helixer gene models, and final model selection was based on support from the protein datasets and LiftOff projections. The result of this was the final set of genome annotations for each genome accession.

Functional annotation was performed for the final set of models using eifunannot v1.5.0 [33], which integrates InterProScan [34] and AHRD [35].

### Assessment of genome annotation models

The quality of genome annotations was assessed via the **ΔBUSCO** measurement (equation 1). This measurement subtracts the number of missing BUSCO genes identified in the genome annotation (PM) from the number of missing BUSCO genes in the genome (GM). The genomic BUSCO genes from the aforementioned hypocreales_odb10.2019-11-20 dataset were identified using both the MetaEuk [29] and compleasm [25] algorithms where the annotated BUSCO genes for the BUSCO protein assessment were extracted from the genomes and original annotation files using gffread v0.12.7. The compleasm algorithm has been shown to be more accurate at detecting BUSCOs than the MetaEuk algorithm in all test cases except distant homologs [25]. As such, we used both algorithms to assess annotation completeness, with compleasm acting as the higher annotation completeness threshold due to increased sensitivity.

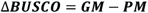

A **ΔBUSCO** score of 0 means that the same number of BUSCO genes are found in the genome and the annotation file. A score below 0 would indicate that there are a greater number of BUSCO genes to be found in the genome than were captured by the annotation process, potentially indicating an issue with the sensitivity of the annotation approach. A score greater than 0 would indicate that the genome annotation captured a greater number of BUSCO genes than was captured by the annotation algorithms employed by BUSCO, which is as expected given additional datasets are normally used to inform annotation. Importantly, a score above 0 is not indicative of any genome annotation quality concern.

Structural similatrity i.e. the comparison of exon-intron chains between annotations was assessed via Mikado compare [36]. Gene-level precision was calculated from pairwise comparisons of the native annotation from each genome (either the original or the final annotation from this study) to the projected annotation from each of the other genomes.

### Assessment of genome assembly quality

Genome assembly metadata was downloaded via ncbi datasets and processed using Rstudio Version 2024.12.0+467. The genome assembly Ilysp1 was excluded from this analysis as it is not deposited on ncbi and so does not have the same assembly statistics available.

### Replicating orthogroup analyses

To assess the impact of computational errors in previous comparative analyses using the original *Fusarium* annotations, we replicated orthogroup analyses by Hill et al. [18] using the same taxa from the reannotated dataset (n=62; Supplementary Table S1). We used OrthoFinder v2.4.0 [37], the same version as used by Hill et al. [18], to infer orthogroups for the reannotated dataset. We performed a phylogenetically informed permutational analysis of variance (PERMANOVA) [38], which used R packages vegan v2.6-4 [39] and RVAideMemoire v0.9-81-2 [40]. Plots were produced in R v4.3.1 [41] using the packages ape v5.7-1 [42], tidyverse v2.0.0 [43], ggtree v3.9.1 [44], ggstats v0.6.0 [45], cowplot v1.1.3 [46], phytools v2.1-2 [47] and ggh4x v0.2.8 [48].

## RESULTS AND DISCUSSION

The reannotation process resulted in a more complete and consistent set of genome annotations across *Fusarium* and allied genera. For the original annotations, the initial **ΔBUSCO** scores, our metric for annotation completeness relative to the assembly, ranged from −684–102 with an average of 8.86 for MetaEuk and −783–6 with an average of −79 for compleasm (Supplementary Fig. S1). The number accessions with a **ΔBUSCO** score below 0 was 17 for MetaEuk and 74 for compleasm. The standard deviation of 109 for MetaEuk and 111 for compleasm showed a wide dispersion of the **ΔBUSCO** scores around the mean, reflecting the inconsistency of annotation completeness across the original annotations. There were notable outliers with particularly poor **ΔBUSCO** scores; accessions GCA_000350365.1 (*Fusarium odoratissimum* Foc4_1.0), GCA_022627095.1 (*Fusarium* sp. 836490-20), GCA_022627115.1 (*Fusarium annulatum* 880149-04), and to a lesser extent GCA_003033665.1 (*Fusarium culmorum* PV) and GCA_022627125.1 (*Fusarium chuoi* 836515-16), indicating that these annotations miss a particularly large number of BUSCO genes that are actually present in the genome assembly.

Comparing the original annotations with the final reannotations, gene annotation completeness was improved for every accession (Fig. 2a, Supplementary Fig. S1), with a final average **ΔBUSCO** score of 91, an average increase of 83 for both MetaEuk and compleasm algorithms. The number of annotations with a **ΔBUSCO** score below 0 was 0 for the MetaEuk algorithm and 11 for compleasm (an improvement from 17 and 74 in the original annotation). An investigation into these 11 accessions showed that the corresponding assemblies all had on average a higher number of contigs as well as a much higher contig N50 and L50 than those accessions with **ΔBUSCO** scores above 0, indicating that these assemblies are more fragmented. Despite this all these accessions benefitted from our reannotation approach with **ΔBUSCO** score improvements ranging from 21-743.

**Figure 2.**
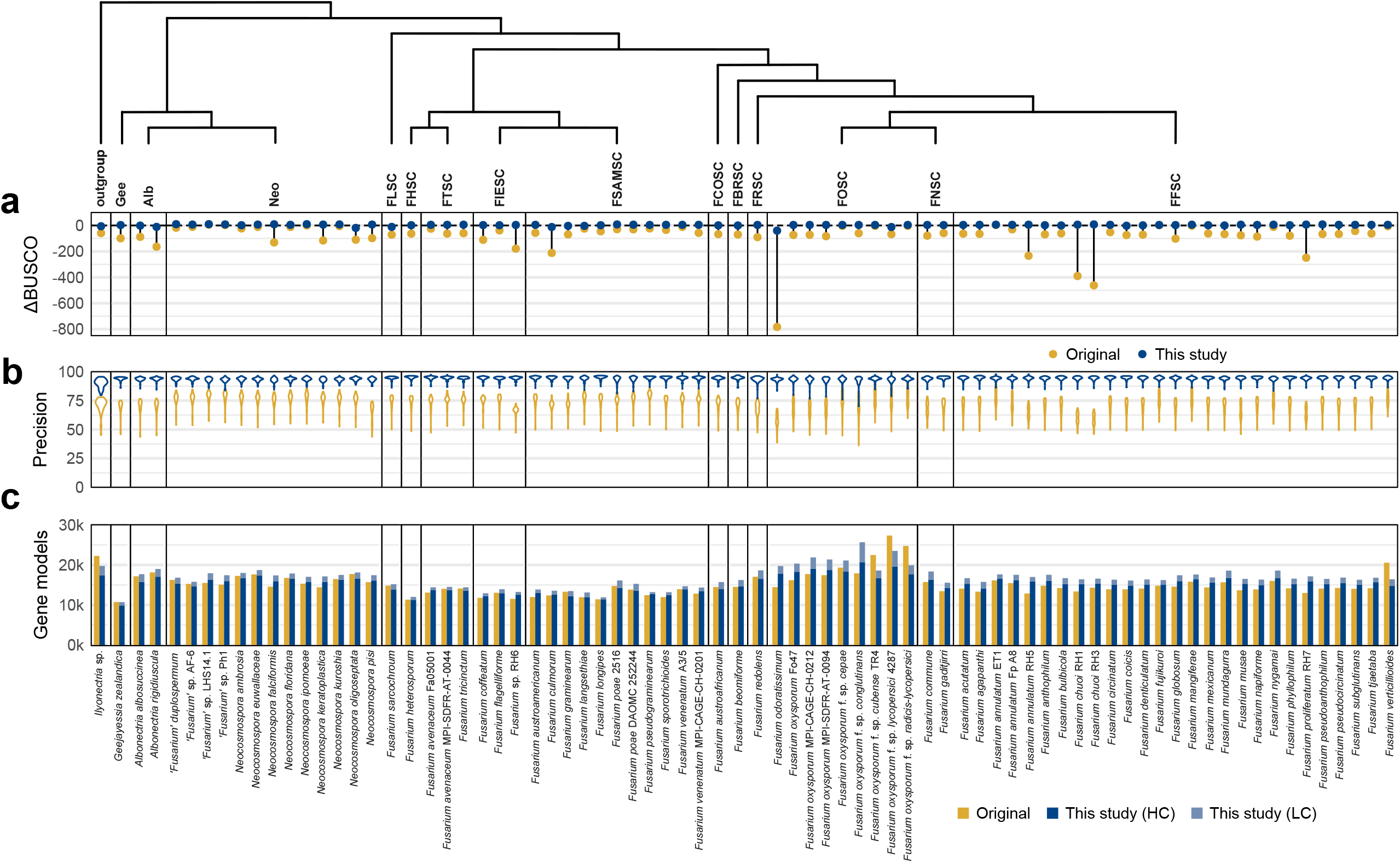
Comparison of the genome annotations before (yellow) and after (blue) reannotation. Species complexes and allied genera are grouped and ordered in accordance with phylogenetic relationships (see Fig. 3a). (a) The difference in BUSCOs missing from the genome assembly and BUSCOs missing from the annotation (**ΔBUSCO**), calculated with compleasm. See Supplementary Fig. S1 for additional comparisons of **ΔBUSCO** when using the Metaeuk versus compleasm algorithms and Supplementary Datasheet S1 for the underlying values. (b) Gene-level precision for exact matches (base F1 of 1) as measured by Mikado compare which indicates how similar each gene model is to the same models across each accession. (c) The total number of gene models, with the reannotations split into high-confidence (HC) and low-confidence (LC) gene models.

The largest gains in gene annotation completeness stemmed from the first round of our reannotation pipeline, which featured a complementary set of *de novo* and homology-based gene annotation methods (Fig. 1). After the first round of the reannotation process the **ΔBUSCO** scores increased by an average of 81 for both MetaEuk and compleasm, with MetaEuk then placing all accessions’ **ΔBUSCO** scores in the positive range. This contrasted with compleasm, which still placed 14 accessions within the negative score range, although each of these accessions still showed a significant improvement with the minimum score having been improved from −783 to −41. The collective consistency of the annotations was markedly improved, with a considerably reduced range of **ΔBUSCO** scores, from 58–146 for MetaEuk and −41–10 for compleasm. The scores were also more centred around the mean, with a standard deviation of 11 and 7 for MetaEuk and compleasm, respectively. The previously highlighted poorly scoring accessions showed major improvements, most marked of which was GCA_000350365.1 (*Fusarium odoratissimum* Foc4_1.0) which showed an improvement from −684 to 58 for Metaeuk and −783 to −41 for Compleasm. This reflected the recovery of 742 BUSCO genes which were previously absent from the genome annotation.

The second round of the reannotation process, which featured a pairwise liftover process serving as a final annotation-unifying step, showed more modest improvements with a **ΔBUSCO** score increase of less than 1 from the first round for both MetaEuk and compleasm. Additionally, both ranges also improved by 1 and the standard deviation remained static indicating no shifts in distributions of **ΔBUSCO** scores across the ranges. However, the second round of reannotation resulted in a further two accessions with **ΔBUSCO** scores above 0 for compleasm, leaving only 12 with a **ΔBUSCO** score of 0 where originally this was 72 (Supplementary Fig. S1).

In addition to annotation completeness, the individual and collective gene content was also far less varied for the final reannotations, with gene-level precision, as measured by Mikado compare, being individually higher and collectively less-varied (Fig. 2b). This measure captures the consistency of intron-exon structure for each gene across the accessions. A high, and less varied gene-level precision indicates that a greater number of gene structures (i.e. exon-intron chains) are identical across the accessions.

Furthermore the number of missing genes, as measured by OMArk [26], was reduced across the accessions (Supplementary Fig. S2). Similar to BUSCO assessment, OMArk estimates completeness based on the proportion of expected conserved ancestral genes present, but OMArk includes conserved multicopy genes (in addition to 1 to 1 orthologues) and so the set of genes that can be assessed is larger. OMArk also evaluates taxonomic consistency by measuring the proportion of protein sequences assigned to known gene families within the same lineage. This approach allows OMArk to assess proteome quality by not only identifying expected proteins but also detecting unexpected (unknown) elements, such as potential contamination or dubious proteins. [26]. Our reannotation process resulted in greater uniformity but also increased the number of gene models classified as being of unknown origin (Supplementary Fig. S2). It is possible that the inconsistent and unknown gene models are indeed erroneous, however their inclusion in the high confidence gene model set indicates that they have >=90% query and target coverage against the public *Fusarium* protein dataset utilised in this study. As such it is also possible that these gene models are valid for the *Fusarium* genus and that the underlying OMArk database used for classification is somewhat incomplete in its repertoire of gene models for this taxonomic group. The reannotation process did remove all gene models flagged as contamination by OMArk, which occurred for a single accession.

The final reannotations resulted in an average increase from 15,162 to 15,277 genes per genome, or 16,751 when including low-confidence gene models (Fig. 2c). While reannotation mostly increased the total number of gene models per genome, in a few cases there was an overall reduction, namely for *F. oxysporum* f. sp. *lycopersici* 4287 (GCA_000149955.2), *F. oxysporum* f. sp. *radicis-lycopersici* (GCA_000260155.3), *Fusarium oxysporum* f. sp. *cubense* TR4 (GCA_000260195.2) and *F. verticillioides* (GCA_000149555.1). These were notably amongst the oldest annotations in the original dataset, perhaps suggesting that the annotation methods used at the time inflated gene model artefacts.

Another merit of our reannotation is that every accession now has more nuanced data associated with gene model predictions, including confidence ratings, UTRs (Supplementary Fig. S3) and predicted isoforms (Supplementary Fig. S4). Where only 27 accessions had any annotated 3’ UTRs and 29 accessions had any 5’ UTRs all accessions are now annotated with both 5’ and 3’ UTRs. Similarly, only 4 accessions had any alternative transcripts in their original annotated (with a mean of 1.24 transcripts per gene) where now all accessions have alternative transcripts (with a mean of 2.1 transcripts per gene).

Genome assemblies and annotations for certain biological groupings, such as the various fungal lifestyles in this dataset, are often produced at the same time for a specific project or initiative, or due to a research group’s specialisation (e.g. [49–51]). This means that differences in genome content observed between groupings may stem from the methods used to generate the data, rather than biology itself. To assess the impact of reannotation on addressing biological questions posed by Hill et al. [18] using the original dataset, we replicated orthogroup presence/absence analyses using the new reannotated dataset. Regarding phylogenomic reconstruction, the species tree produced using STAG was virtually unchanged (Fig. 3a). There were two bipartitions in the tree which differed, but one was notably in a difficult-to-resolve clade that is already known to exhibit a lot of incongruence [18] and associated branches had low support values in the trees produced both before and after reannotation. Our results suggest that phylogenomic reconstruction may be relatively robust to genome annotation inconsistencies at the magnitude present in the original dataset. As many phylogenomic approaches typically use a heavily filtered set of single-copy orthogroups with a high taxon occupancy, it is possible that issues regarding genome annotation are often circumvented in phylogenomic analyses. However, while topology was largely unaffected, improving annotation consistency did have the merit of increasing the proportion of well supported branches (from 56% in Hill et al. to 61% in this study).

**Figure 3.**
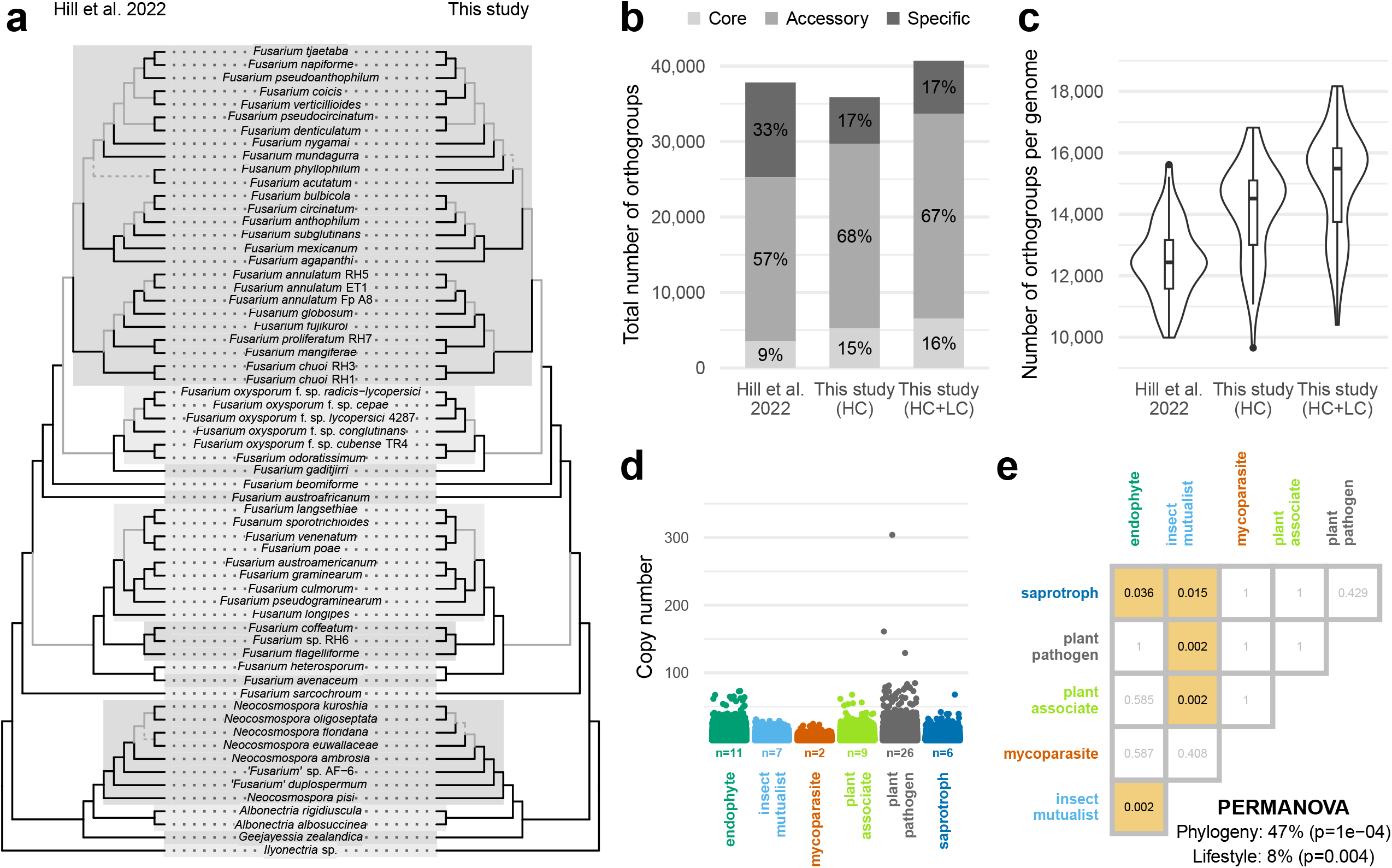
Replicating analyses on 62 *Fusarium* taxa by Hill et al. [18] using the reannotated dataset produced in this study. HC=high-confidence gene models, LC=low-confidence gene models. (a) Tanglegram comparing total-evidence species trees generated with STAG. Dashed branches indicate contradictory topology. Grey branches indicate fewer than 1/3 of gene trees supporting the branch. *Fusarium* species complexes are indicated with alternated highlights. (b) Total number of orthogroups recovered by OrthoFinder, with colour distinguishing whether orthogroups were core, accessory or specific within fusarioid taxa. See Supplementary Fig. S5 for the breakdown of core, accessory and specific orthogroups for each individual genome. (c) Number of orthogroups per genome. (d) Variation in copy number across all (HC+LC) orthogroups for different lifestyles. Points are jittered and the number of strains for each lifestyle is reported under x-axis labels. (e) Pairwise PERMANOVA results indicating whether all orthogroup (HC+LC) content was significantly different between lifestyles. Coloured boxes indicate significant p-values (<0.05). Global PERMANOVA results are reported at the bottom.

Reannotation resulted in an increased proportion of core and accessory orthogroups, and a reduction in taxon-specific orthogroups (Fig. 3b), consistent with the finding that the number of species or strain specific genes is often inflated by mixed bioinformatics approaches [13]. The average number of orthogroups per genome increased by 13.2% (or 20.8% if including both HC and LC gene models; Fig. 3c). The reannotated dataset agreed with the findings of Hill et al. [18] that the genomes of plant pathogenic taxa contained copy-number outliers (Fig. 3d) but the difference when compared to other lifestyles including endophytes was not as strongly marked as in the original analysis. The reannotated dataset also supported previous results that insect mutualist fungi had significantly different orthogroup content compared to most other lifestyles (Fig. 3e), however the descriptive power of phylogeny was increased substantially (from 35% to 47% in this study) and the power of lifestyle marginally reduced (from 9% to 8% in this study) presumably due to the recovery of artefactually missing gene models. We also found a significant difference in orthogroup content between endophyte and insect mutualist genomes, which was previously only found for a subset of genes predicted to be candidate secreted effector proteins (CSEPs) [18]. This suggests that the noise created by annotation errors may also mask potentially meaningful biological signal.

From a biological perspective the biggest caveat to our approach is that it only focused on protein-coding genes. Incorporation of the projection, *de novo* prediction and validation of non-coding RNAs, as was performed for protein-coding genes, is a non-trivial venture and was outside of the scope of this work. An additional biological caveat is that the success of this method is likely to be closely tied to the evolutionary distance, or genome similarity, between the genomes. We can see this when we plot the number of transferred genes as a function of evolutionary distance for a single example (Supplementary Fig. S6). The dependence of alignment-based annotation performance upon sequence similarity has been previously discussed and demonstrated by [22].

From a computational perspective the major limitation is the scalability of this pan-annotation approach. The method used herein required multiple iterations of pairwise comparisons of gene models for each possible pair of genomes. Given a larger set of genomes, or larger genomes with an increased gene count, and the computational resources required are likely to be significant. Future development of this work to surpass the pairwise comparison problem is currently underway. Another limitation is that this approach naturally cannot rectify errors/missing genes originating from low assembly quality [52], and so there should still be caution in interpreting comparative results if there is doubt as to the quality of the original genome sequencing or assembly of accessions.

Comparative analyses are fundamental for understanding biology, and it is natural that researchers should want to use existing data for these purposes. New strategies to revise and update annotations must be developed to combat annotation errors and prevent the propagation of computational artefacts across research. We have integrated existing bioinformatics tools into a new workflow centred on pairwise liftovers and demonstrated an improvement in genome annotation quality both individually and as a collective. We used this approach to improve existing annotations, but it could also be used to efficiently fill annotation gaps – as of writing NCBI comprises ∼18k fungal genomes but only ∼5k have an associated annotation. Even in new genome sequencing projects where the same annotation methodology is used across all genomes, a pairwise liftover approach can be integrated into the annotation workflow to improve consistency (e.g. [53]). We hope this new pan-annotation resource for *Fusarium* will be of use to the fungal research community.

## Supporting information

Supplementary Datasheet S1

## ACKNOWLEDGEMENTS

We thank the attendees of the quarterly meetings for the Earlham Institute’s strategic research programme Decoding Biodiversity, as well as Mark McMullan, for valuable discussion and feedback.

## DATA AVAILABILITY

All reannotations are deposited in Zenodo doi:10.5281/zenodo.13829922. Workflow code and sample commands are available from https://github.com/EI-CoreBioinformatics/FusariumPanAnno.

## REFERENCES

1. Heather JM, Chain B. The sequence of sequencers: The history of sequencing DNA. Genomics 2016; 107:1–8. 10.1016/j.ygeno.2015.11.003.

2. Duan H, Jones AW, Hewitt T, et al. Physical separation of haplotypes in dikaryons allows benchmarking of phasing accuracy in Nanopore and HiFi assemblies with Hi-C data. Genome Biol 2022; 23:84. 10.1186/s13059-022-02658-2.

3. The Darwin Tree of Life Project Consortium. Sequence locally, think globally: The Darwin Tree of Life Project. PNAS 2022; 119:e2115642118. 10.1073/pnas.2115642118.

4. Mc Cartney AM, Formenti G, Mouton A, et al. The European Reference Genome Atlas: piloting a decentralised approach to equitable biodiversity genomics. npj biodivers 2024; 3:28. 10.1038/s44185-024-00054-6.

5. Guigó R. Genome annotation: From human genetics to biodiversity genomics. Cell Genomics 2023; 3:100375. 10.1016/j.xgen.2023.100375.

6. Hamese S, Mugwanda K, Takundwa M, et al. Recent advances in genome annotation and synthetic biology for the development of microbial chassis. Journal of Genetic Engineering and Biotechnology 2023; 21:156. 10.1186/s43141-023-00598-3.

7. Ceballos G, Ehrlich PR, Barnosky AD, et al. Accelerated modern human–induced species losses: Entering the sixth mass extinction. Sci. Adv. 2015; 1:e1400253. 10.1126/sciadv.1400253.

8. Berger B, Yu YW. Navigating bottlenecks and trade-offs in genomic data analysis. Nat Rev Genet 2023; 24:235–250. 10.1038/s41576-022-00551-z.

9. Scalzitti N, Jeannin-Girardon A, Collet P, et al. A benchmark study of ab initio gene prediction methods in diverse eukaryotic organisms. BMC Genomics 2020; 21:293. 10.1186/s12864-020-6707-9.

10. Dimonaco NJ, Aubrey W, Kenobi K, et al. No one tool to rule them all: prokaryotic gene prediction tool annotations are highly dependent on the organism of study. Bioinformatics 2022; 38:1198–1207. 10.1093/bioinformatics/btab827.

11. Siezen RJ, van Hijum SAFT. Genome (re-)annotation and open-source annotation pipelines. Microbial Biotechnology 2010; 3:362–369. 10.1111/j.1751-7915.2010.00191.x.

12. Klimke W, O’Donovan C, White O, et al. Solving the Problem: Genome Annotation Standards before the Data Deluge. Stand Genomic Sci 2011; 5:168–193. 10.4056/sigs.2084864.

13. Weisman CM, Murray AW, Eddy SR. Mixing genome annotation methods in a comparative analysis inflates the apparent number of lineage-specific genes. Curr Biol 2022; 32:2632–2639. 10.1016/j.cub.2022.04.085.

14. Weisman CM, Murray AW, Eddy SR. Many, but not all, lineage-specific genes can be explained by homology detection failure. PLoS Biol 2020; 18:e3000862. 10.1371/journal.pbio.3000862.

15. Prada CF, Boore JL. Gene annotation errors are common in the mammalian mitochondrial genomes database. BMC Genomics 2019; 20:73. 10.1186/s12864-019-5447-1.

16. Monnahan PJ, Michno J-M, O’Connor C, et al. Using multiple reference genomes to identify and resolve annotation inconsistencies. BMC Genomics 2020; 21:281. 10.1186/s12864-020-6696-8.

17. Kress A, Poch O, Lecompte O, et al. Real or fake? Measuring the impact of protein annotation errors on estimates of domain gain and loss events. Front Bioinform 2023; 3:1178926. 10.3389/fbinf.2023.1178926.

18. Hill R, Buggs RJA, Vu DT, et al. Lifestyle Transitions in Fusarioid Fungi are Frequent and Lack Clear Genomic Signatures. Mol Biol Evol 2022; 39:msac085. 10.1093/molbev/msac085.

19. Armer VJ, Kroll E, Darino M, et al. Navigating the Fusarium species complex: Host-range plasticity and genome variations. Fungal Biology 2024; 10.1016/j.funbio.2024.07.004.

20. van Dam P, Fokkens L, Schmidt SM, et al. Effector profiles distinguish formae speciales of Fusarium oxysporum. Environmental Microbiology 2016; 18:4087–4102. 10.1111/1462-2920.13445.

21. Czislowski E, Zeil-Rolfe I, Aitken EAB. Effector profiles of endophytic Fusarium associated with asymptomatic banana (Musa sp.) hosts. International Journal of Molecular Sciences 2021; 22:2508. 10.3390/ijms22052508.

22. Brůna T, Li H, Guhlin J, et al. Galba: genome annotation with miniprot and AUGUSTUS. BMC Bioinformatics 2023; 24:327. 10.1186/s12859-023-05449-z.

23. Stiehler F, Steinborn M, Scholz S, et al. Helixer: cross-species gene annotation of large eukaryotic genomes using deep learning. Bioinformatics 2021; 36:5291–5298. 10.1093/bioinformatics/btaa1044.

24. Shumate A, Salzberg SL. Liftoff: accurate mapping of gene annotations. Bioinformatics 2021; 37:1639–1643. 10.1093/bioinformatics/btaa1016.

25. Huang N, Li H. compleasm: a faster and more accurate reimplementation of BUSCO. Bioinformatics 2023; 39:btad595. 10.1093/bioinformatics/btad595.

26. Nevers Y, Warwick Vesztrocy A, Rossier V, et al. Quality assessment of gene repertoire annotations with OMArk. Nat Biotechnol 2025; 43:124–133. 10.1038/s41587-024-02147-w.

27. Pertea G, Pertea M. GFF Utilities: GffRead and GffCompare. F1000Res 2020; 9:304. 10.12688/f1000research.23297.2.

28. EI-CoreBioinformatics. minos - a gene model consolidation pipeline for genome annotation projects. 2023; https://github.com/EI-CoreBioinformatics/minos.

29. Manni M, Berkeley MR, Seppey M, et al. BUSCO Update: Novel and Streamlined Workflows along with Broader and Deeper Phylogenetic Coverage for Scoring of Eukaryotic, Prokaryotic, and Viral Genomes. Mol Biol Evol 2021; 38:4647–4654. 10.1093/molbev/msab199.

30. Kaithakottil GG, Swarbreck D. EIRepeat - EI Repeat Identification Pipeline. 2024; https://github.com/EI-CoreBioinformatics/eirepeat.

31. Smit A, Hubley R. RepeatModeler Open-1.0. 2015; http://www.repeatmasker.org.

32. Smit A, Hubley R, Green P. RepeatMasker Open-4.0. 2015; http://www.repeatmasker.org.

33. Kaithakottil G, Swarbreck D. eifunannot. 2024; https://github.com/EI-CoreBioinformatics/eifunannot.

34. Jones P, Binns D, Chang HY, et al. InterProScan 5: genome-scale protein function classification. Bioinformatics 2014; 30:1236–1240. 10.1093/bioinformatics/btu031.

35. Hallab A, Klee K, Boecker F, et al. Automated Assignment of Human Readable Descriptions (AHRD). 2023; https://github.com/groupschoof/AHRD.

36. Venturini L, Caim S, Kaithakottil GG, et al. Leveraging multiple transcriptome assembly methods for improved gene structure annotation. GigaScience 2018; 7: 10.1093/gigascience/giy093.

37. Emms DM, Kelly S. OrthoFinder: Phylogenetic orthology inference for comparative genomics. Genome Biol 2019; 20:238. 10.1101/466201.

38. Mesny F, Vannier N. Detecting the effect of biological categories on genome composition. 2020; https://github.com/fantin-mesny/Effect-Of-Biological-Categories-On-Genomes-Composition.

39. Oksanen J, Blanchet FG, Friendly M, et al. vegan: Community Ecology Package. 2019; https://cran.r-project.org/web/packages/vegan/index.html.

40. Hervé M. RVAideMemoire: Testing and Plotting Procedures for Biostatistics. 2020; https://cran.r-project.org/package=RVAideMemoire.

41. R Core Team. R: A language and environment for statistical computing. 2023; https://www.r-project.org/.

42. Paradis E, Schliep K. Ape 5.0: An environment for modern phylogenetics and evolutionary analyses in R. Bioinformatics 2019; 35:526–528. 10.1093/bioinformatics/bty633.

43. Wickham H, Averick M, Bryan J, et al. Welcome to the Tidyverse. J Open Source Softw 2019; 4:1686. 10.21105/joss.01686.

44. Yu G, Smith DK, Zhu H, et al. GGTREE: an R package for visualization and annotation of phylogenetic trees with their covariates and other associated data. Methods Ecol Evol 2017; 8:28–36. 10.1111/2041-210X.12628.

45. Larmarange J. ggstats: Extension to ‘ggplot2’ for Plotting Stats. 2024; https://larmarange.github.io/ggstats/.

46. Wilke CO. cowplot: Streamlined Plot Theme and Plot Annotations for ‘ggplot2’. 2024; https://cran.r-project.org/package=cowplot.

47. Revell LJ. phytools 2.0: an updated R ecosystem for phylogenetic comparative methods (and other things). PeerJ 2024; 12:e16505. 10.7717/peerj.16505.

48. van den Brand T. ggh4x: Hacks for ‘ggplot2’. 2024; https://CRAN.R-project.org/package=ggh4x.

49. Martino E, Morin E, Grelet GA, et al. Comparative genomics and transcriptomics depict ericoid mycorrhizal fungi as versatile saprotrophs and plant mutualists. New Phytologist 2018; 217:1213–1229. 10.1111/nph.14974.

50. Marqués-Gálvez JE, Miyauchi S, Paolocci F, et al. Desert truffle genomes reveal their reproductive modes and new insights into plant–fungal interaction and ectendomycorrhizal lifestyle. New Phytologist 2021; 229:2917–2932. 10.1111/nph.17044.

51. Llewellyn T, Nowell RW, Aptroot A, et al. Metagenomics Shines Light on the Evolution of “Sunscreen” Pigment Metabolism in the Teloschistales (Lichen-Forming Ascomycota). Genome Biol Evol 2023; 15:evad002. 10.1093/gbe/evad002.

52. Denton JF, Lugo-Martinez J, Tucker AE, et al. Extensive Error in the Number of Genes Inferred from Draft Genome Assemblies. PLoS Computational Biology 2014; 10:e1003998.

53. Hill R, Grey M, Olivera Fedi M, et al. Evolutionary genomics reveals variation in structure and genetic content implicated in virulence and lifestyle in the genus Gaeumannomyces. BMC Genom 2024; in press.

