## Supplementary Material for "Leveraging existing data to maximise quality and consistency across gene model annotations: a *Fusarium* pan-annotation"

**Supplementary Table S1** Summary of the genomes used in this study. Asterisks (*) indicate accessions for new additions that weren't used in comparative analyses by Hill et al. [1]. Dagger (†) indicates duplicate RefSeq accessions. HC=high-confidence, LC=low-confidence.

| Species | Strain | Accession | Original annotation reference | Number of genes Original annotation | Number of genes This study [Total(HC;LC)] |
|---|---|---|---|---|---|
| *Fusarium verticillioides* | NRRL 20956 | GCA_000149555.1 | [2] | 20,553 | 16,406(14,751;1,655) |
| *Fusarium oxysporum* f. sp. *lycopersici* | 4287 | GCA_000149955.2 | [2] | 27,347 | 23,499(19,605;3,894) |
| *† *Fusarium oxysporum* f. sp. *lycopersici* | 4287 | GCF_000149955.1 | [2] | 27,346 | 23,447(19,489;3,958) |
| *Neocosmospora pisi* | NRRL 44580; 77-13-4 | GCA_000151355.1 | [3] | 15,708 | 17,439(16,097;1,342) |
| *Fusarium graminearum* | NRRL 31084; PH-1 | GCA_000240135.3 | [4] | 13,313 | 13,474(12,187;1,287) |
| *Fusarium oxysporum* f. sp. *radicis-lycopersici* | 26381 | GCA_000260155.3 | [5] | 24,739 | 19,925(17,705;2,220) |
| *Fusarium oxysporum* f. sp. *cubense* TR4 | NRRL 54006 | GCA_000260195.2 | [5] | 22,487 | 18,616(16,704;1,912) |
| *Fusarium pseudograminearum* | CS3096 | GCA_000303195.2 | [6] | 12,447 | 13,211(12,736;475) |
| *† *Fusarium pseudograminearum* | CS3096 | GCF_000303195.2 | [6] | 12,397 | 13,209(12,734;475) |
| *Fusarium odoratissimum* | Foc4_1.0 | GCA_000350365.1 | [7] | 14,459 | 19,741(17,841;1,900) |
| *Fusarium avenaceum* | Fa05001 | GCA_000769215.1 | [8] | 13,092 | 14,424(13,754;670) |
| *Fusarium langsethiae* | Fl201059; 9821-16-1 | GCA_001292635.1 | [9] | 11,940 | 13,121(11,891;1,230) |
| *Fusarium agapanthi* | NRRL 31653 | GCA_001654555.2 | [10] | 13,310 | 15,765(14,089;1,676) |
| *Fusarium poae* | 2516 | GCA_001675295.1 | [11] | 14,740 | 16,124(14,236;1,888) |
| *Fusarium nygamai* | CS10214 | GCA_002894225.1 | | 16,019 | 18,598(16,682;1,916) |
| *Fusarium beomiforme* | NRRL 25174 | GCA_002980475.2 | [10] | 14,521 | 16,231(14,692;1,539) |
| *Fusarium longipes* | NRRL 20695 | GCA_003012285.1 | [12] | 11,421 | 11,921(11,347;574) |
| *Fusarium flagelliforme* | NRRL 13405 | GCA_003012295.1 | [12] | 13,046 | 13,935(12,943;992) |
| *Fusarium sporotrichioides* | NRRL 3299 | GCA_003012315.1 | [12] | 11,961 | 13,196(12,443;753) |
| *Fusarium culmorum* | PV | GCA_003033665.1 | [13] | 12,378 | 13,556(12,599;957) |
| *Fusarium coffeatum* | FIESC_28 | GCA_003316985.1 | | 11,779 | 12,932(12,216;716) |
| *Fusarium oxysporum* f. sp. *cepae* | FoC_Fus2 | GCA_003615085.1 | [14] | 19,342 | 21,105(18,316;2,789) |
| *Fusarium annulatum* | Fp_A8 | GCA_003615215.1 | [14] | 15,449 | 17,526(16,210;1,316) |
| *Neocosmospora kuroshia* | UCR3666 | GCA_003698175.1 | | 16,485 | 17,501(16,308;1,193) |
| *'Fusarium' duplospermum* | NRRL 62584; AF-8 | GCA_003946985.1 | [10] | 16,263 | 16,818(15,345;1,473) |

| Species | Strain | Accession | Original annotation reference | Number of genes Original annotation | Number of genes This study [Total(HC;LC)] |
|---|---|---|---|---|---|
| *Neocosmospora oligoseptata* | NRRL 62579; AF-4 | GCA_003946995.1 | [10] | 17,740 | 18,115(16,545;1,570) |
| *Neocosmospora floridana* | NRRL 62606 | GCA_003947005.1 | [10] | 16,762 | 17,861(16,606;1,255) |
| *'Fusarium'* sp. AF-6 | NRRL 62590 | GCA_003947015.1 | [10] | 15,274 | 15,780(14,639;1,141) |
| *Neocosmospora ambrosia* | NRRL 20438 | GCA_003947045.1 | [10] | 17,262 | 17,998(16,683;1,315) |
| *Neocosmospora euwallaceae* | UCR1854 | GCA_003957675.1 | | 17,630 | 18,726(17,339;1,387) |
| *Albonectria albosuccinea* | NRRL 20459 | GCA_012931995.1 | [10] | 17,169 | 17,687(15,765;1,922) |
| *Fusarium acutatum* | NRRL 13308 | GCA_012932015.1 | [10] | 14,081 | 16,696(15,286;1,410) |
| *Fusarium austroafricanum* | NRRL 53441 | GCA_012932025.1 | [10] | 14,502 | 15,725(14,016;1,709) |
| *Fusarium gaditjirri* | NRRL 45417; FRC M-8754 | GCA_013266175.1 | [10] | 13,454 | 15,557(14,279;1,278) |
| *Fusarium sarcochroum* | NRRL 20472 | GCA_013266185.1 | [10] | 14,827 | 15,199(13,893;1,306) |
| *Geejayessia zealandica* | NRRL 22465 | GCA_013266195.1 | [10] | 10,733 | 10,717(9,828;889) |
| *Albonectria rigidiuscula* | NRRL 13412 | GCA_013266205.1 | [10] | 18,126 | 18,984(17,058;1,926) |
| *Fusarium anthophilum* | NRRL 25214 | GCA_013364935.1 | [10] | 14,797 | 17,538(16,042;1,496) |
| *Fusarium austroamericanum* | NRRL 2903 | GCA_013364965.1 | [10] | 11,984 | 13,865(12,996;869) |
| *Fusarium pseudoanthophilum* | NRRL 25211 | GCA_013395995.1 | [10] | 14,088 | 16,499(15,219;1,280) |
| *Fusarium napiforme* | NRRL 25196 | GCA_013396005.1 | [10] | 13,896 | 16,339(14,983;1,356) |
| *Fusarium mexicanum* | NRRL 53147 | GCA_013396015.1 | [10] | 14,387 | 16,884(15,633;1,251) |
| *Fusarium phyllophilum* | NRRL 13617 | GCA_013396025.1 | [10] | 14,160 | 16,578(15,240;1,338) |
| *Fusarium pseudocircinatum* | NRRL 36939 | GCA_013396035.1 | [10] | 14,263 | 16,856(15,496;1,360) |
| *Fusarium subglutinans* | NRRL 66333 | GCA_013396075.1 | [10] | 14,040 | 16,295(15,070;1,225) |
| *Fusarium globosum* | NRRL 26131 | GCA_013396165.1 | [10] | 14,590 | 17,423(15,954;1,469) |
| *Fusarium denticulatum* | NRRL 25311 | GCA_013396175.1 | [10] | 14,097 | 16,369(15,111;1,258) |
| *Fusarium circinatum* | NRRL 25331 | GCA_013396185.1 | [10] | 13,905 | 16,316(15,017;1,299) |
| *Fusarium tjaetaba* | NRRL 66243 | GCA_013396195.1 | [10] | 14,181 | 16,802(15,686;1,116) |
| *† *Fusarium tjaetaba* | NRRL 66243 | GCF_013396195.1 | [10] | 14,181 | 16,802(15,687;1,115) |
| *Fusarium mundagurra* | NRRL 66235 | GCA_013396205.1 | [10] | 15,666 | 18,590(16,758;1,832) |
| *Fusarium heterosporum* | NRRL 20693 | GCA_013396295.1 | [10] | 11,332 | 12,007(11,289;718) |

| Species | Strain | Accession | Original annotation reference | Number of genes Original annotation | Number of genes This study [Total(HC;LC)] |
|---|---|---|---|---|---|
| *Fusarium bulbicola* | NRRL 25176 | GCA_013758895.1 | [10] | 14,231 | 16,706(15,228;1,478) |
| *Fusarium coicis* | NRRL 66233 | GCA_013781345.1 | [10] | 13,908 | 16,100(14,854;1,246) |
| *Fusarium oxysporum* f. sp. *conglutinans* | Fo5176 | GCA_014154955.1 | [15] | 17,912 | 25,671(20,694;4,977) |
| * *Fusarium avenaceum* | MPI-SDFR-AT-0044 | GCA_020744115.1 | [16] | 14,042 | 14,513(13,723;790) |
| * *Fusarium venenatum* | MPI-CAGE-CH-0201 | GCA_020744135.1 | [16] | 12,845 | 14,367(13,472;895) |
| * *Fusarium* commune | MPI-SDFR-AT-0072 | GCA_020744335.1 | [16] | 15,730 | 18,365(16,287;2,078) |
| * *Fusarium oxysporum* | MPI-CAGE-CH-0212 | GCA_020744355.1 | [16] | 17,726 | 21,889(19,045;2,844) |
| * *Fusarium oxysporum* | MPI-SDFR-AT-0094 | GCA_020744455.1 | [16] | 17,444 | 21,366(18,714;2,652) |
| * *Fusarium tricinctum* | MPI-SDFR-AT-0068 | GCA_020744515.1 | [16] | 14,106 | 14,421(13,574;847) |
| *Fusarium* sp. | RH6; 836490-20 | GCA_022627095.1 | [1] | 11,533 | 13,258(12,550;708) |
| *Fusarium chuoi* | RH3; 836445-12-1 | GCA_022627105.1 | [1] | 14,313 | 16,584(15,361;1,223) |
| *Fusarium annulatum* | RH5; 880149-04 | GCA_022627115.1 | [1] | 12,880 | 16,993(15,874;1,119) |
| *Fusarium chuoi* | RH1; 836515-16 | GCA_022627125.1 | [1] | 13,380 | 16,414(15,210;1,204) |
| *Fusarium proliferatum* | RH7; 836489-13 | GCA_022627135.1 | [1] | 13,009 | 17,158(15,927;1,231) |
| * Neocosmospora ipomoeae (=*Fusarium solani-melongenae)* | CRI 24-3 | GCA_023101225.1 | [17] | 15,320 | 17,027(15,785;1,242) |
| * ‘*Fusarium’* sp. | Ph1 | GCA_025433565.1 | [18] | 15,084 | 17,434(15,960;1,474) |
| * ‘*Fusarium’* sp. | LHS14.1 | GCA_025433615.1 | [18] | 15,522 | 17,929(16,285;1,644) |
| *Fusarium venenatum* | A3/5 | GCA_900007375.1 | [19] | 13,946 | 14,682(13,905;777) |
| *† *Fusarium venenatum* | A3/5 | GCF_900007375.1 | [19] | 13,932 | 14,678(13,908;770) |
| *Fusarium mangiferae* | MRC7560 | GCA_900044065.1 | [20] | 15,804 | 17,610(16,421;1,189) |
| *† *Fusarium mangiferae* | MRC7560 | GCF_900044065.1 | [20] | 15,797 | 17,609(16,419;1,190) |
| *Fusarium annulatum* | ET1 | GCA_900067095.1 | [21] | 16,143 | 17,640(16,534;1,106) |
| *† *Fusarium annulatum* | ET1 | GCF_900067095.1 | [21] | 16,136 | 17,640(16,533;1,107) |
| * *Fusarium oxysporum* | Fo47 | GCF_013085055.1 | [22] | 16,202 | 20,324(18,287;2,037) |
| * *Fusarium poae* | DAOMC 252244 | GCF_019609905.1 | [23] | 13,851 | 15,273(13,546;1,727) |
| * *Fusarium musae* | F31 | GCF_019915245.1 | [24] | 13,672 | 16,514(15,281;1,233) |
| * *Fusarium redolens* | MPI-CAGE-AT-0023 | GCF_020744475.1 | [16] | 17,049 | 18,649(16,515;2,134) |

| Species | Strain | Accession | Original annotation reference | Number of genes Original annotation | Number of genes This study [Total(HC;LC)] |
|---|---|---|---|---|---|
| * *Neocosmospora keratoplastica (=Fusarium keratoplasticum)* | Fu6.1 | GCF_025433545.1 | [18] | 14,423 | 17,143(15,766;1,377) |
| * *Neocosmospora falciformis (=Fusarium falciforme)* | Fu3.1 | GCF_026873545.1 | [18] | 14,574 | 17,383(15,951;1,432) |
| *Fusarium fujikuroi* | IMI 58289 | GCF_900079805.1 | [25] | 14,810 | 16,220(15,017;1,203) |
| *Ilyonectria* sp. | Ilysp1 | Ilysp1 | [26] | 22,250 | 19,742(17,411;2,331) |


1. Hill R, Buggs RJA, Vu DT, et al. Lifestyle Transitions in Fusarioid Fungi are Frequent and Lack Clear Genomic Signatures. *Mol Biol Evol* 2022; 39:msac085. https://doi.org/10.1093/molbev/msac085.
2. Ma LJ, van der Does HC, Borkovich KA, et al. Comparative genomics reveals mobile pathogenicity chromosomes in *Fusarium*. *Nature* 2010; 464:367–373. https://doi.org/10.1038/nature08850.
3. Coleman JJ, Rounsley SD, Rodriguez-Carres M, et al. The genome of *Nectria haematococca*: Contribution of supernumerary chromosomes to gene expansion. *PLoS Genetics* 2009; 5:e1000618. https://doi.org/10.1371/journal.pgen.1000618.
4. Cuomo CA, Güldener U, Xu J, et al. The *Fusarium graminearum* Genome Reveals a Link Between Localized Polymorphism and Pathogen Specialization. *Science* 2007; 317:1400–1403. https://doi.org/10.1126/science.1143708.
5. Delulio GA, Guo L, Zhang Y, et al. Kinome Expansion in the *Fusarium oxysporum* Species Complex Driven by Accessory Chromosomes. *mSphere* 2018; 3:e00231-18.
6. Gardiner DM, McDonald MC, Covarelli L, et al. Comparative Pathogenomics Reveals Horizontally Acquired Novel Virulence Genes in Fungi Infecting Cereal Hosts. *PLoS Pathogens* 2012; 8:e1002952. https://doi.org/10.1371/journal.ppat.1002952.
7. Guo L, Han L, Yang L, et al. Genome and transcriptome analysis of the fungal pathogen *Fusarium oxysporum* f. sp. *cubense* causing banana vascular wilt disease. *PLoS ONE* 2014; 9:e95543. https://doi.org/10.1371/journal.pone.0095543.
8. Lysøe E, Harris LJ, Walkowiak S, et al. The Genome of the Generalist Plant Pathogen *Fusarium avenaceum* Is Enriched with Genes Involved in Redox, Signaling and Secondary Metabolism. *PLoS ONE* 2014; 9:e112703. https://doi.org/10.1371/journal.pone.0112703.
9. Lysøe E, Frandsen RJN, Divon HH, et al. Draft genome sequence and chemical profiling of *Fusarium langsethiae*, an emerging producer of type A trichothecenes. *International Journal of Food Microbiology* 2016; 221:29–36. https://doi.org/10.1016/j.ijfoodmicro.2016.01.008.
10. Kim H-S, Lohmar JM, Busman M, et al. Identification and distribution of gene clusters required for synthesis of sphingolipid metabolism inhibitors in diverse species of the filamentous fungus *Fusarium*. *BMC Genomics* 2020; 21:510. https://doi.org/10.1186/s12864-020-06896-1.
11. Vanheule A, Audenaert K, Warris S, et al. Living apart together: crosstalk between the core and supernumerary genomes in a fungal plant pathogen. *BMC Genomics* 2016; 17:670. https://doi.org/10.1186/s12864-016-2941-6.
12. Proctor RH, McCormick SP, Kim HS, et al. Evolution of structural diversity of trichothecenes, a family of toxins produced by plant pathogenic and entomopathogenic fungi. *PLoS Pathogens* 2018; 14:e1006946. https://doi.org/10.1371/journal.ppat.1006946.

13. Schmidt R, Durling MB, de Jager V, et al. Deciphering the genome and secondary metabolome of the plant pathogen *Fusarium culmorum*. *FEMS Microbiology Ecology* 2018; 94:fiy078. https://doi.org/10.1093/femsec/fiy078.
14. Armitage AD, Taylor A, Sobczyk MK, et al. Characterisation of pathogen-specific regions and novel effector candidates in *Fusarium oxysporum* f. sp. *cepae*. *Scientific Reports* 2018; 8:13530. https://doi.org/10.1038/s41598-018-30335-7.
15. Fokkens L, Guo L, Dora S, et al. A Chromosome-Scale Genome Assembly for the *Fusarium oxysporum* Strain Fo5176 To Establish a Model *Arabidopsis*-Fungal Pathosystem. *G3: Genes Genom Genet* 2020; 10:3549–3555. https://doi.org/10.25387/G3.12707897.
16. Mesny F, Miyauchi S, Thiergart T, et al. Genetic determinants of endophytism in the *Arabidopsis* root mycobiome. *Nat Commun* 2021; 12:7227. https://doi.org/10.1038/s41467-021-27479-y.
17. Xie S-Y, Ma T, Zhao N, et al. Whole-Genome Sequencing and Comparative Genome Analysis of *Fusarium solani-melongenae* Causing Fusarium Root and Stem Rot in Sweetpotatoes. *Microbiol Spectr* 2022; 10:e00683-22. https://doi.org/10.1128/spectrum.00683-22.
18. Hoh DZ, Lee H-H, Wada N, et al. Comparative genomic and transcriptomic analyses of trans-kingdom pathogen *Fusarium solani* species complex reveal degrees of compartmentalization. *BMC Biol* 2022; 20:236. https://doi.org/10.1186/s12915-022-01436-7.
19. King R, Brown NA, Urban M, et al. Inter-genome comparison of the Quorn fungus *Fusarium venenatum* and the closely related plant infecting pathogen *Fusarium graminearum*. *BMC Genomics* 2018; 19:269. https://doi.org/10.1186/s12864-018-4612-2.
20. Britz H, Steenkamp ET, Coutinho TA, et al. Two new species of *Fusarium* section *Liseola* associated with mango malformation. *Mycologia* 2002; 94:722–730. https://doi.org/10.1080/15572536.2003.11833199.
21. Niehaus EM, Münsterkötter M, Proctor RH, et al. Comparative 'Omics' of the *Fusarium fujikuroi* Species Complex Highlights Differences in Genetic Potential and Metabolite Synthesis. *Genome Biology and Evolution* 2016; 8:3574–3599. https://doi.org/10.1093/gbe/evw259.
22. Wang B, Yu H, Jia Y, et al. Chromosome-Scale Genome Assembly of *Fusarium oxysporum* Strain Fo47, a Fungal Endophyte and Biocontrol Agent. *MPMI* 2020; 33:1108–1111. https://doi.org/10.1094/MPMI-05-20-0116-A.
23. Witte TE, Harris LJ, Nguyen HDT, et al. Apicidin biosynthesis is linked to accessory chromosomes in *Fusarium poae* isolates. *BMC Genomics* 2021; 22:591. https://doi.org/10.1186/s12864-021-07617-y.
24. Degradi L, Tava V, Kunova A, et al. Telomere to Telomere Genome Assembly of *Fusarium musae* F31, Causal Agent of Crown Rot Disease of Banana. *MPMI* 2021; 34:1455–1457. https://doi.org/10.1094/MPMI-05-21-0127-A.
25. Wiemann P, Sieber CMK, von Bargen KW, et al. Deciphering the Cryptic Genome: Genome-wide Analyses of the Rice Pathogen *Fusarium fujikuroi* Reveal Complex Regulation of Secondary Metabolism and Novel Metabolites. *PLoS Pathogens* 2013; 9:e1003475. https://doi.org/10.1371/journal.ppat.1003475.
26. Liao HL, Bonito G, Rojas JA, et al. Fungal endophytes of *Populus trichocarpa* alter host phenotype, gene expression, and rhizobiome composition. *MPMI* 2019; 32:853–864. https://doi.org/10.1094/MPMI-05-18-0133-R.

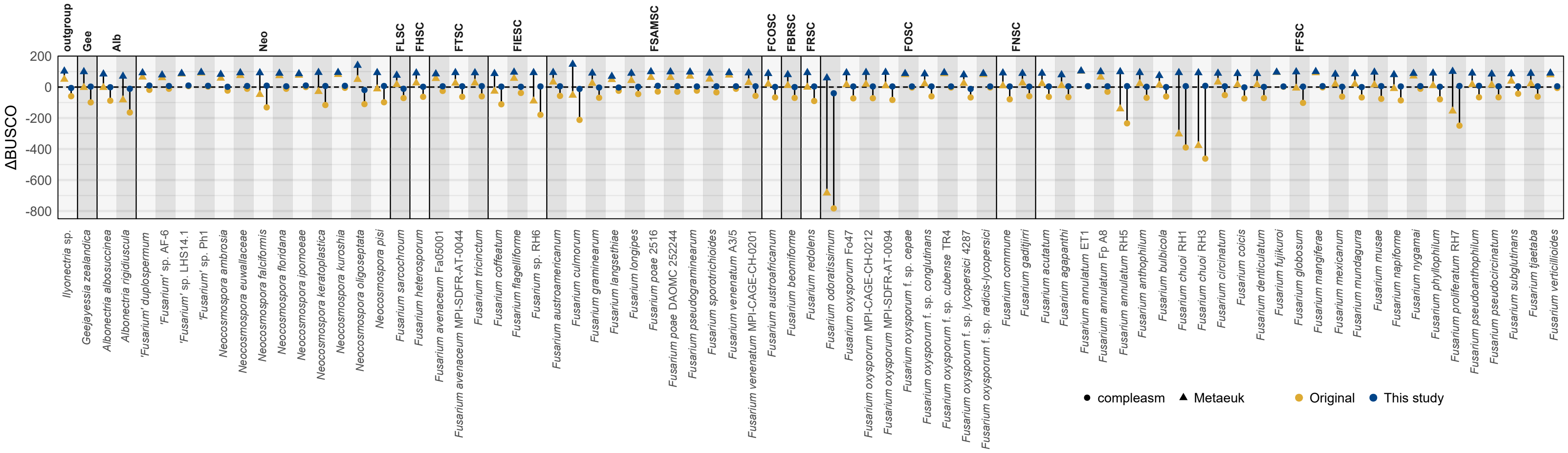


**Supplementary Fig. S1** The difference in BUSCOs missing from the genome assembly and BUSCOs missing from the annotation (∆BUSCO). Shapes distinguish results produced using the Metaeuk algorithm versus the compleasm algorithm.

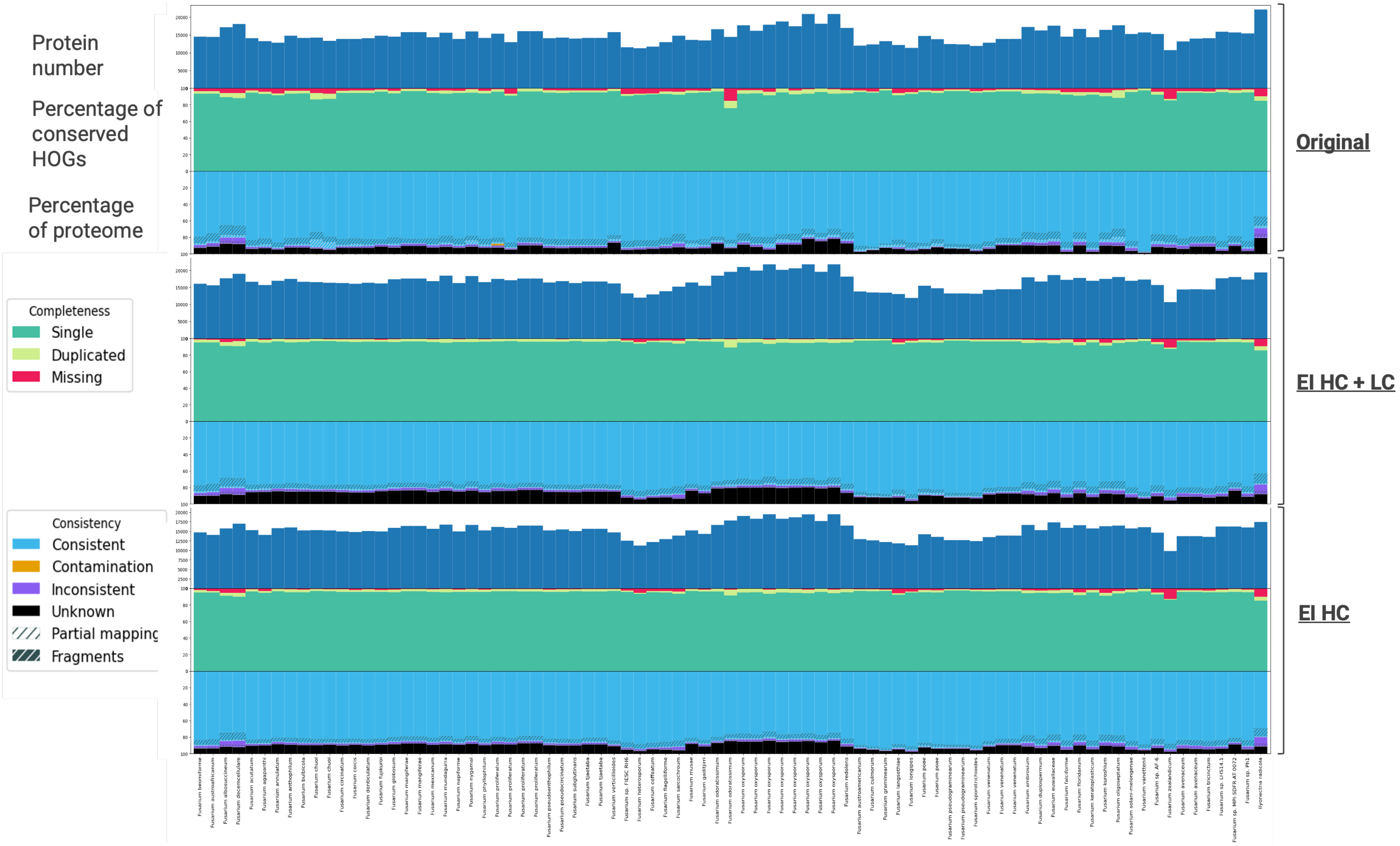


**Supplementary Fig. S2** Completeness and consistency of conserved genes as determined by OMArk for the original annotations versus the reannotations in this study (HC=high-confidence; LC=low-confidence).

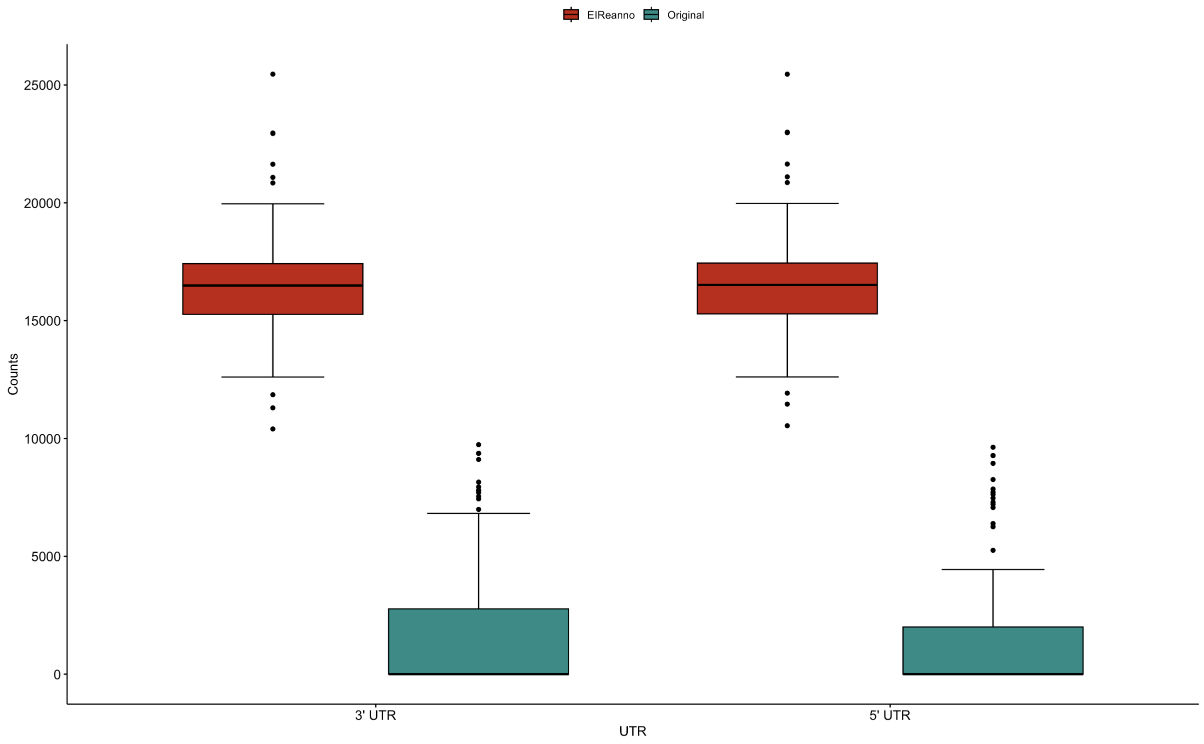


**Supplementary Fig. S3** Number of UTR annotations per genome.

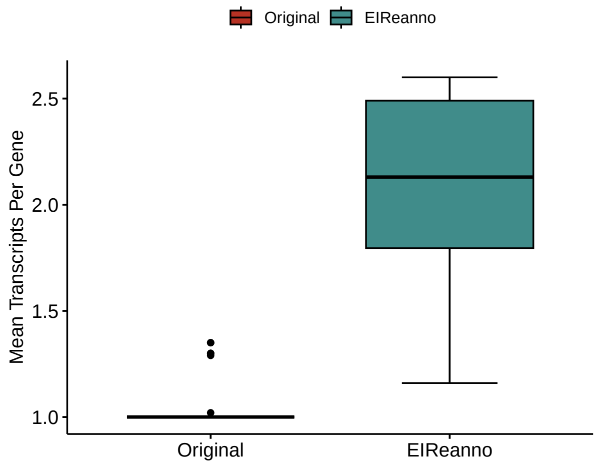


**Supplementary Fig. S4** Number of transcripts per gene.

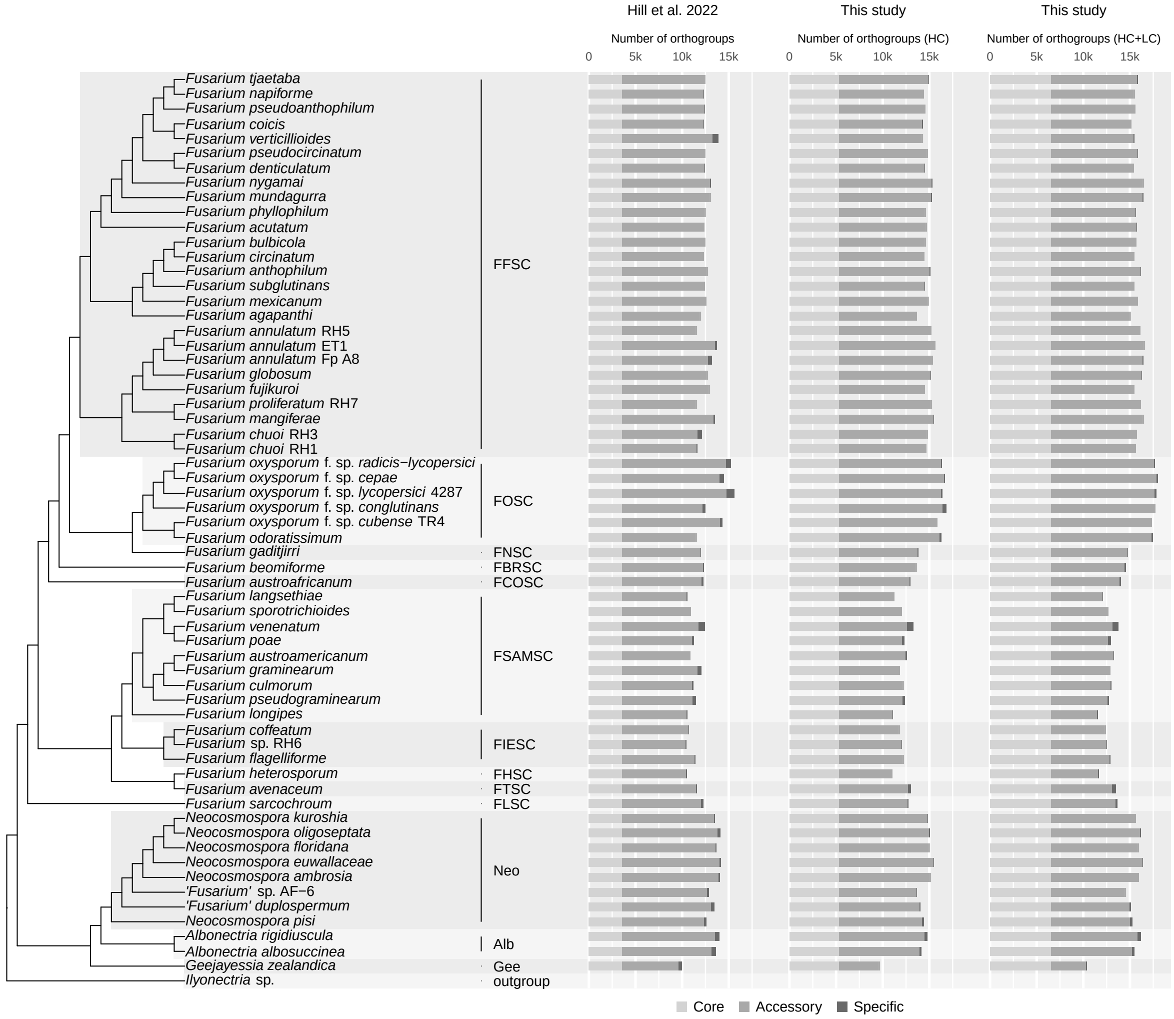


**Supplementary Fig. S5** Number of orthogroups recovered by OrthoFinder for the original dataset from Hill et al. (2022) and the reannotated dataset from this study, with colour distinguishing whether orthogroups were core, accessory or specific within fusarioid taxa. HC=high-confidence gene models, LC=low-confidence gene models.

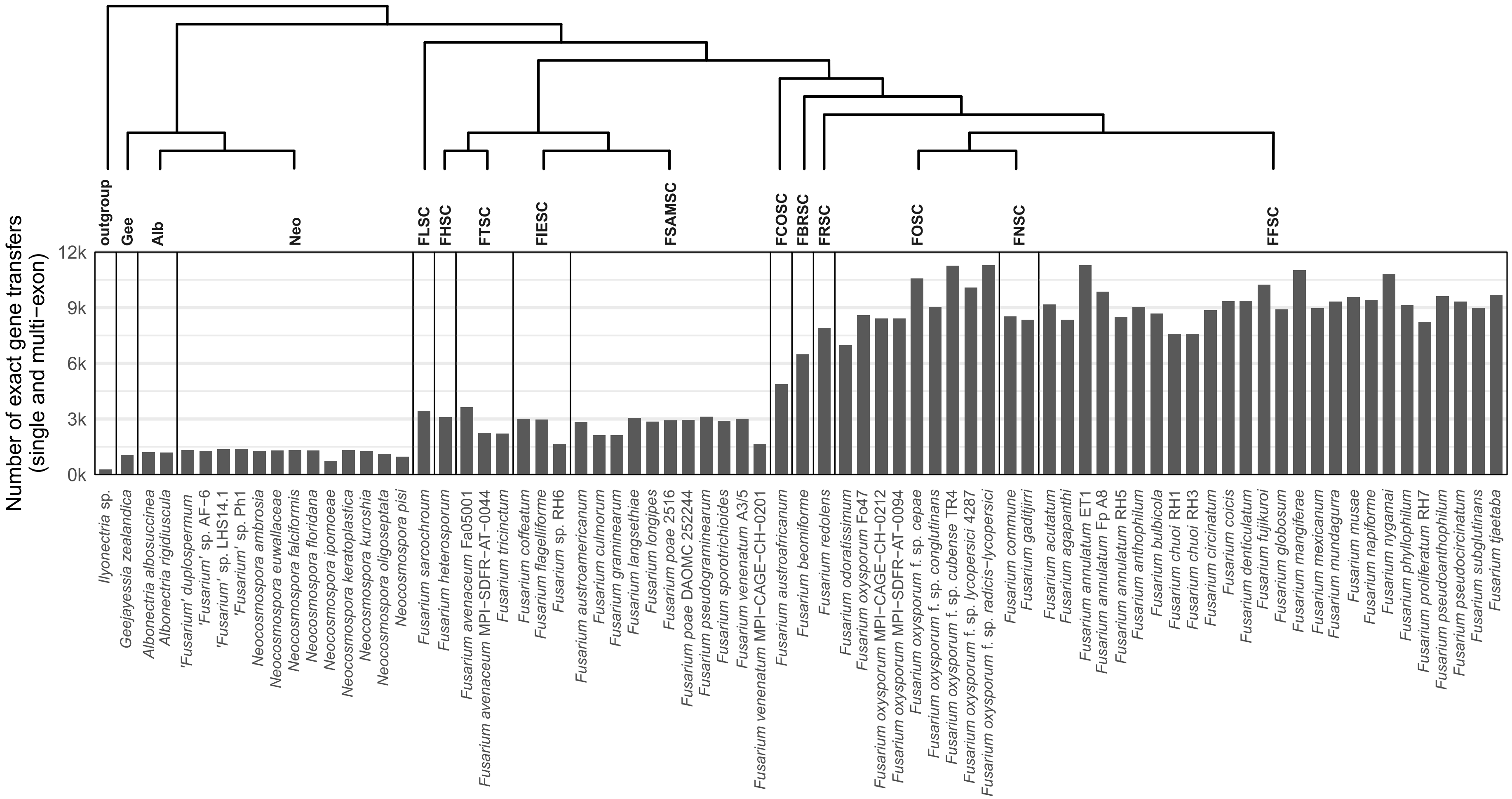


**Supplementary Fig. S6** The number of exact gene LiftOvers for GCA_000149555.1 (*Fusarium verticillioides* NRRL 20956, FFSC).
